# High-resolution structure determination of sub-100 kilodalton complexes using conventional cryo-EM

**DOI:** 10.1101/489898

**Authors:** Mark A. Herzik, Mengyu Wu, Gabriel C. Lander

## Abstract

Determining high-resolution structures of biological macromolecules with masses of less than 100 kilodaltons (kDa) has long been a goal of the cryo-electron microscopy (cryo-EM) community. While the Volta Phase Plate has enabled cryo-EM structure determination of biological specimens of this size range, use of this instrumentation is not yet fully automated and can present technical challenges. Here, we show that conventional defocus-based cryo-EM methodologies can be used to determine the high-resolution structures of specimens amassing less than 100 kDa using a transmission electron microscope operating at 200 keV coupled with a direct electron detector. Our ∼2.9 Å structure of alcohol dehydrogenase (82 kDa) proves that bound ligands can be resolved with high fidelity, indicating that these methodologies can be used to investigate the molecular details of drug-target interactions. Our ∼2.8 Å and ∼3.2 Å resolution structures of methemoglobin demonstrate that distinct conformational states can be identified within a dataset for proteins as small as 64 kDa. Furthermore, we provide the first sub-nanometer cryo-EM structure of a protein smaller than 50 kDa.

---

In recent years, technical advances in cryo-electron microscopy (cryo-EM) single-particle analysis (SPA) have propelled the technique towards the forefront of structural biology, enabling the direct visualization of biological macromolecules in near-native states at increasingly higher resolutions^1^. Notably, cryo-EM enables three-dimensional (3D) structure determination of biological specimens in a vitrified state without the requirement of crystallization^2^, which has not only greatly increased the throughput of high-resolution structure determination, but has also allowed for the 3D visualization of macromolecular complexes previously deemed intractable for structural studies due to size, conformational heterogeneity, and/or compositional variability^3-5^. Indeed, determining ∼3 Å resolution reconstructions of stable, conformationally and/or compositionally homogeneous specimens by SPA has become almost routine, with an increasing number of structures at ∼2 Å resolution or better now being reported^6-8^. This resolution regime has also expanded the potential of cryo-EM SPA for structure-based drug design, particularly for targets that are less amenable to other structure-determination techniques due to limited sample quantity or recalcitrance to crystallization.

Despite these advances, the radiation sensitivity of ice-embedded proteins and the low signal-to-noise ratio (SNR) of cryo-EM images nonetheless impede routine structure determination for all samples, and specimen size remains a considerable limiting factor in SPA cryo-EM. Indeed, although SPA reconstructions of molecules as small as 38 kilodaltons (kDa) have been theorized to be achievable^9^, this feat has yet to be realized. To date, only three macromolecular complexes smaller than 100 kDa have been resolved to high resolution (i.e. better than 4 Å) using SPA cryo-EM: the ∼ 3.8 Å resolution reconstruction of 93 kDa isocitrate dehydrogenase^10^, the ∼3.2 Å resolution reconstruction of 64 kDa methemoglobin (mHb)^5^, and the ∼3.2 Å resolution reconstruction of 52 kDa streptavidin^11^. Due to the limited success in imaging smaller macromolecules by cryo-EM, the technique has primarily been used to visualize large complexes, with approximately 99% of all cryo-EM reconstructions resolved to better than 5 Å resolution comprising macromolecules amassing *>*200 kDa.

We previously demonstrated that a transmission electron microscope (TEM) operating at 200 keV equipped with a K2 Summit direct electron detector (DED) could be used to resolve a ∼150 kDa protein complex to ∼2.6 Å using conventional defocus-based SPA methods^12^. Here, we expand upon our previous results and show that biological specimens amassing *<*100 kDa can be resolved to better than 3 Å resolution using similar imaging approaches. The resulting reconstructions possess well-resolved density for bound co-factors, metal ligands, as well as ordered water molecules. We further demonstrate that conformational heterogeneity in specimens of this size range can be discerned. Finally, we provide the first sub-nanometer single-particle cryo-EM structure of a sub-50 kDa macromolecular complex – the 43 kDa catalytic domain of protein kinase A.

## Results and Discussion

Our prior success in resolving the structure of a sub-200 kDa complex to better than 3 Å resolution demonstrated that conventional defocus-based methodologies provided sufficient SNR to confidently assign 3D orientations to biological specimens that were previously thought to be too small to image^12^. This prompted us to see if high-resolution reconstructions of macromolecules *<*100 kDa in size could be achieved with conventional approaches, provided sufficiently thin ice and high particle density could be attained to maximize accuracy of contrast transfer function (CTF) estimation. All specimens in this study were imaged using a base model (i.e. excluding imaging accessories such as a phase plate or energy filter) Talos Arctica TEM equipped with a K2 Summit DED operating in counting mode using the Leginon^13^ automated data collection software. TEM column alignments were performed as described previously^12,14^ with the following modification to maximize parallel illumination: after minimizing the spread of gold powder diffraction rings, the camera length was increased to 5.7 m and both the size of the caustic spot and the diffraction astigmatism were iteratively minimized in order to optimize parallel illumination of the sample (see Methods).

### Structure of 82 kDa alcohol dehydrogenase at ∼2.9 Å resolution

We first decided to target the 82 kDa homodimeric enzyme alcohol dehydrogenase (ADH)^15^, as the structure of this complex had been previously determined to near-atomic resolution (e.g. better than 1.3 Å resolution) by X-ray diffraction^16^, providing an atomic model for validation of our results. Notably, as ADH is a nicotinamide adenine dinucleotide (NADH)-binding protein, part of our motivation for selecting this enzyme for structure determination was to test our ability to resolve the bound NADH cofactor, as this would serve as important proof-of-principle for the utility of SPA for structure-based drug design of similarly sized samples.

We further purified commercially sourced horse liver ADH (Sigma Aldrich) to homogeneity and prepared grids of the frozen-hydrated specimen for cryo-EM data collection (see Methods) (Supplementary Fig. 1). Despite the small size of ADH, multiple views of the homodimer could be clearly distinguished in the aligned images even when using a nominal defocus range of −0.5 *μ*m to −1.6 *μ*m, contrary to previous estimations that significant underfocus would be required to image small (*<*200 kDa) macromolecules using conventional EM^17^ (Supplementary Fig. 2). However, the large disparity in orthogonal dimensions of ADH (∼100 Å versus ∼40 Å, see Fig. 1) posed a challenge during two-dimensional (2D) classification. To overcome this, two successive rounds of RELION^18,19^ reference-free 2D classification were performed, the first using a 110 Å soft circular mask to identify particles comprising the longer side/tilted views, followed by a second round of 2D classification of the remaining data using a 60 Å soft mask to identify particles comprising the “end-on” views (Fig. 1a, Supplementary Fig. 2e). These combined particles were subsequently subjected to several rounds of 3D autorefinement and classification with C2 symmetry enforced to yield a final ∼2.9 Å reconstruction, as determined by gold-standard Fourier shell correlation (FSC) at 0.143^20-22^ (see Methods) (Fig. 1, Supplementary Table 1, and Supplementary Fig. 2). These same particles refined to 3.4 Å resolution without imposing symmetry (Supplementary Fig. 2).

**Figure 1.**
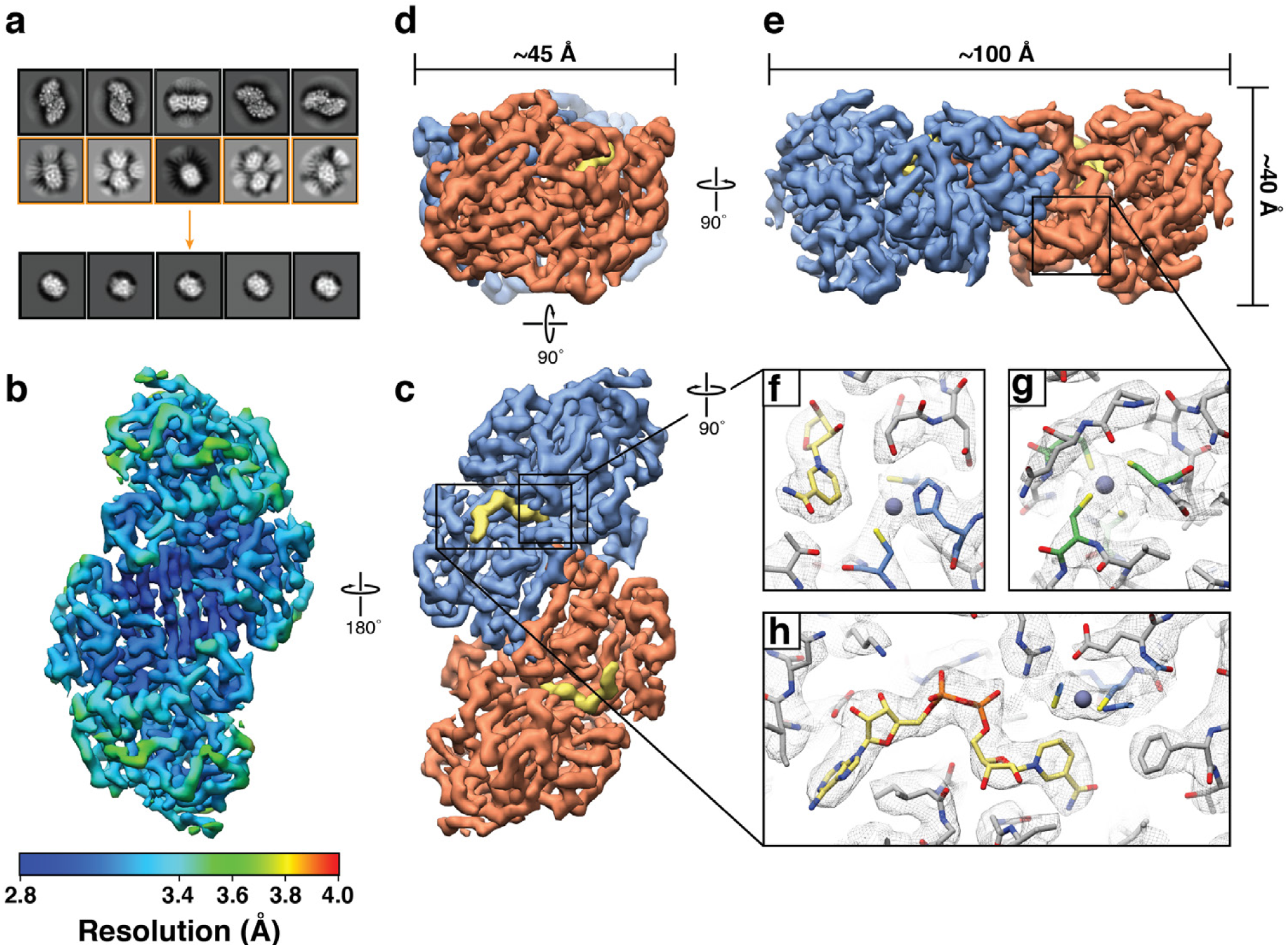
∼2.9 Å resolution cryo-EM reconstruction of 82 kDa horse liver alcohol dehydrogenase. **(a)** Representative reference-free 2D class averages of horse liver alcohol dehydrogenase (ADH). Particles comprising the 2D classes highlighted in yellow were subsequently further classified using a smaller soft circular mask (see methods). **(b)** Final cryo-EM reconstruction colored by estimated local resolution estimated with BSOFT^23^. **(c-e)** Orthogonal views of the ADH EM density (colored by subunit) showing the disparity in particle dimensions. The segmented NADH EM density is shown in yellow. **(f-h)** Zoomed-in views of the EM density (gray mesh) for the ADH active-site zinc, structural zinc site, and active site, respectively. Residues involved in coordinating the active-site zinc (blue), the structural site zinc (green) or interacting with NADH (yellow) are shown in stick representation. The zinc atoms are shown as purple spheres.

Inspection of the map revealed side-chain density for most of the ADH polypeptide with pronounced backbone features for the entire macromolecule. Indeed, local resolution estimates^23^ indicated that the majority of the map was resolved to better than ∼3.4 Å resolution, with the core of ADH resolved to ∼2.8 Å resolution (Fig. 1b). Moreover, there is unambiguous density for the bound NADH co-factor, a catalytic zinc ion within the active site of each protomer, as well as a structural zinc ion coordinated by four cysteine residues (Fig. 1). Importantly, the positions of these ligands are corroborated by the X-ray crystal structure of horse ADH (PDB ID: 2JHF). This reconstruction clearly establishes that *<*100 kDa complexes can be resolved to better than 3 Å resolution using conventional approaches using a 200 keV TEM without the need for a phase plate or energy filter. Furthermore, our ability to confidently identify bound ligands and co-factors within our ADH EM density effectively demonstrates the utility of conventional SPA methods for structure-based drug design and other small molecule research involving similarly sized specimens.

### Structures of distinct human methemoglobin states at ∼2.8 Å and ∼3.2 Å resolution

Khoshouei *et al*. ^5^ recently demonstrated that a TEM operating at 300 keV combined with a Volta phase plate (VPP) and quantum energy filter could be used to resolve human hemoglobin (Hb), a ~64-kDa heterotetrameric heme-containing protein, to ∼3.2 Å resolution. It has been speculated that the increased SNR afforded by the VPP was integral for structure determination of Hb^5^. However, our ability to resolve ADH to high resolution indicated that conventional defocus-based imaging methodologies provided sufficient SNR for structure determination, and that we had not yet reached the size limit of this approach. We therefore sought to determine the structure of Hb using similar strategies. Briefly, vitrified Hb specimens were prepared from lyophilized human Hb (Sigma Aldrich) (see Methods). UV-VIS absorbance measurements of the solubilized Hb sample confirmed that the bound heme was indeed in the ferric (Fe^3+^) oxidation state, hereby referred to as methemoglobin (metHb).

Images of frozen-hydrated metHb were collected similarly as described for ADH (see Methods) (Supplementary Fig. 3). Notably, orthogonal views of metHb were discernible by eye (Fig. 2b), even in micrographs imaged using underfocus values as low as ∼700 nm. RELION reference-free 2D classification yielded detailed class averages exhibiting secondary structural elements and various recognizable views of tetrameric metHb (Fig. 2c). Three parallel 3D classifications were performed to select for unique particles comprising the best-resolved classes across each classification (see Methods, Supplementary Fig. 3). An additional round of no-alignment 3D classification revealed two distinct conformational states of metHb: state 1 (∼2.8 Å resolution) closely matches the “Near R2” state previously described by Shibayama *et al.*^24^ (Cα root-mean-square-deviation (RMSD) 0.4 Å, PDB ID: 4N7P) (Fig. 3a) and state 2 (∼3.2 Å resolution) agrees well with the “Between R and R2” state (Cα RMSD 0.5 Å, PDB ID: 4N7N) (Fig. 3b). Comparison of the two states following superposition of an αβ dimer from each molecule revealed an approximately 7^*°*^ rigid-body rotation of one αβ dimer with respect to the other with the rotation axis centered about the dimer–dimer interface, similar to those movements previously described^24^ (Fig. 3c). These results demonstrate that distinct, biologically relevant conformational states of a sub-100 kDa complex can feasibly be resolved to high-resolution using single-particle cryo-EM.

**Figure 3.**
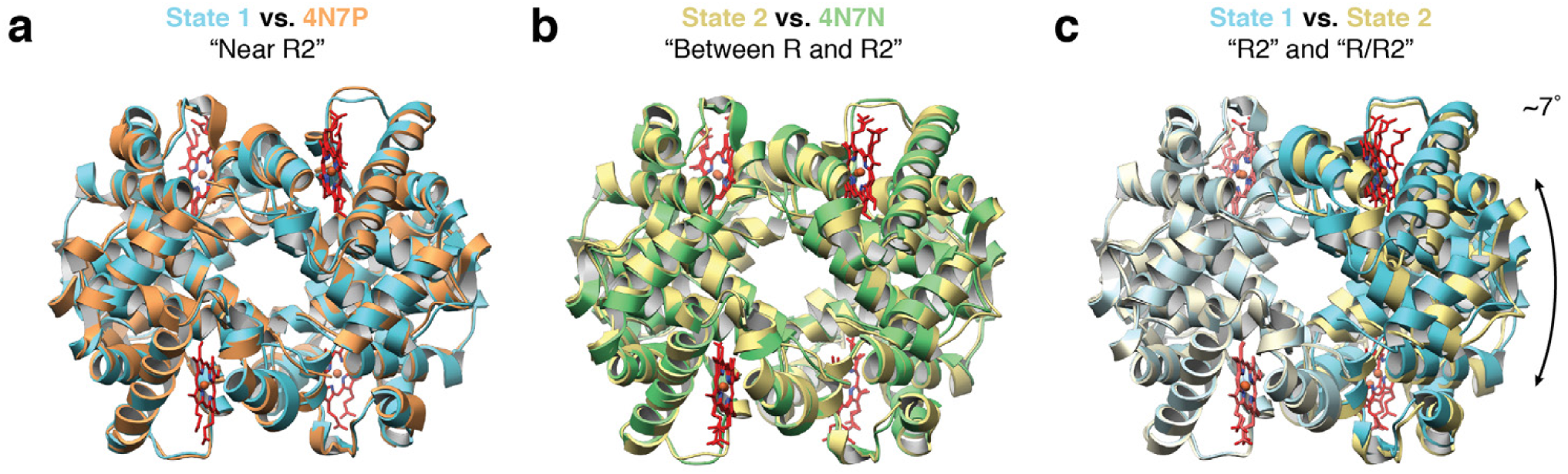
Multiple conformational states of metHb determined using single particle cryo-EM. **(a)** Cartoon representation of metHb state 1 (∼2.8 Å resolution, blue) superposed with PDB ID: 4N7P (chains A-D, orange). **(b)** Cartoon representation of metHb state 2 (∼3.2 Å resolution, yellow) superposed with PDB ID: 4N7N (chains E-H, green). **(c)** Superposing the α1 and β1 subunits of states 1 and 2 emphasizes the difference in between these two conformations as a ∼7^*°*^ rotation of the α2 and β2 subunits.

**Figure 2.**
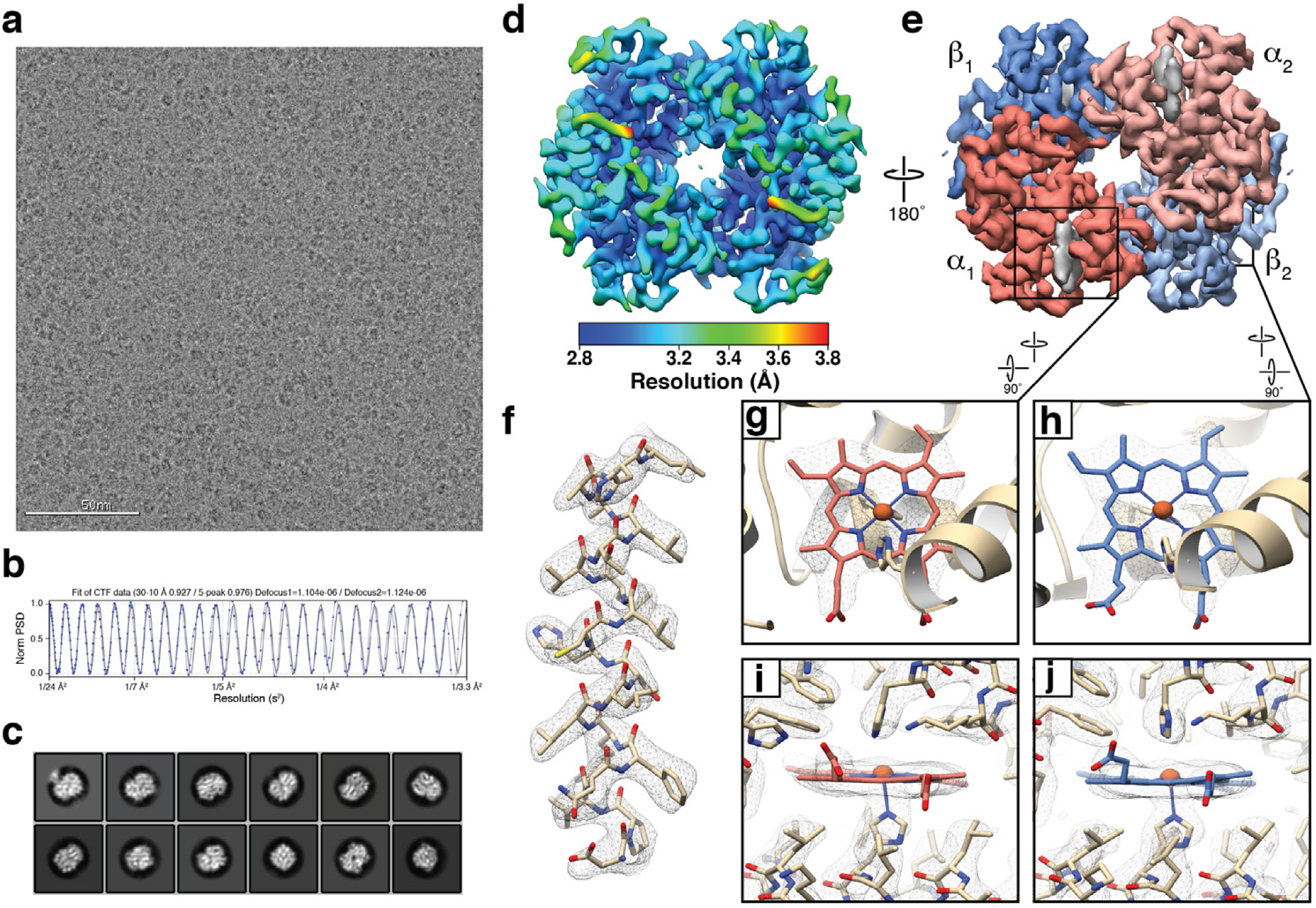
∼2.8 Å resolution cryo-EM reconstruction of ∼64 kDa human methemoglobin. **(a)** Representative motion-corrected electron micrograph of human methemoglobin (metHb) embedded in vitreous ice recorded at ∼1.1 *μ*m defocus (scale bar, 50 nm). **(b)** 1-dimensional plot of the contrast transfer function (CTF) Thon rings (black line) and the CTF estimated with CTFFIND4^33^ (blue line). **(c)** Representative reference-free 2D class averages showing secondary structure elements. **(d-e)** Final ∼2.8 Å resolution metHb cryo-EM density colored by local resolution (estimated using BSOFT^23^)and subunit with the segmented EM density for the heme cofactors colored gray, respectively. **(f)** EM density (gray mesh) zoned 2 Å around an α-helix comprising residues 94-113 from the α subunit. **(g-j)** EM density in the vicinity of the heme cofactors from subunit α1 and β2. Lower panels are 90^*°*^ rotations of upper panels. Residues are shown in stick representation (colored wheat) and the heme cofactors are colored according to the subunit coloring in (e). The heme iron atoms are shown as orange spheres.

Local resolution estimates indicated that most of both maps were resolved to better than ∼3.5 Å, which is consistent with our ability to resolve side-chain densities throughout the map (Fig. 2f). Notably, the heme pockets are estimated to be the best-resolved regions in both states (Supplementary Fig. 3c,e), and the quality of the heme densities in both α and β subunits of state 1 enabled us to confidently discern the location of the vinyl groups extending from the porphyrin ring to unambiguously assign the orientation of the heme moiety in each pocket (Fig. 2e). Furthermore, the well-resolved regions of state 1 contain putative density for ordered water molecules (Supplementary Fig. 3h) that are conserved between those observed in a previously obtained crystal structure (PDB ID: 2DN1). Our ability to discern subtle conformational differences in a target as small as 64 kDa effectively underscores the utility of this approach in examining the conformational dynamics of similarly small biological systems.

### Towards a high-resolution structure of the ∼43 kDa catalytic subunit of protein kinase A

Processing of the metHb dataset required two successive rounds of 2D classification, partially due to the presence of “contaminating” ∼32-kDa αβ heterodimers, which surprisingly accounted for ∼20% of the data (Supplementary Fig. 3). Subsequent 2D classification of these particles yielded class averages with defined secondary structural elements (Fig. 4). Furthermore, 2D projections of a simulated EM density generated using an αβ dimer from PDB ID: 4N7N exhibited striking similarity to these class averages (Fig. 4). However, generation of a 3D reconstruction was unsuccessful, likely due to preferred orientation and/or poor signal-to-noise ratio of the particles in vitreous ice. Despite this, our results for the metHb tetramer indicated that we had not yet reached the size limit resolvable by our imaging methodologies, and the promising 2D averages of the metHb dimers prompted us to continue investigating the lower molecular weight limit of cryo-EM SPA.

**Figure 4.**
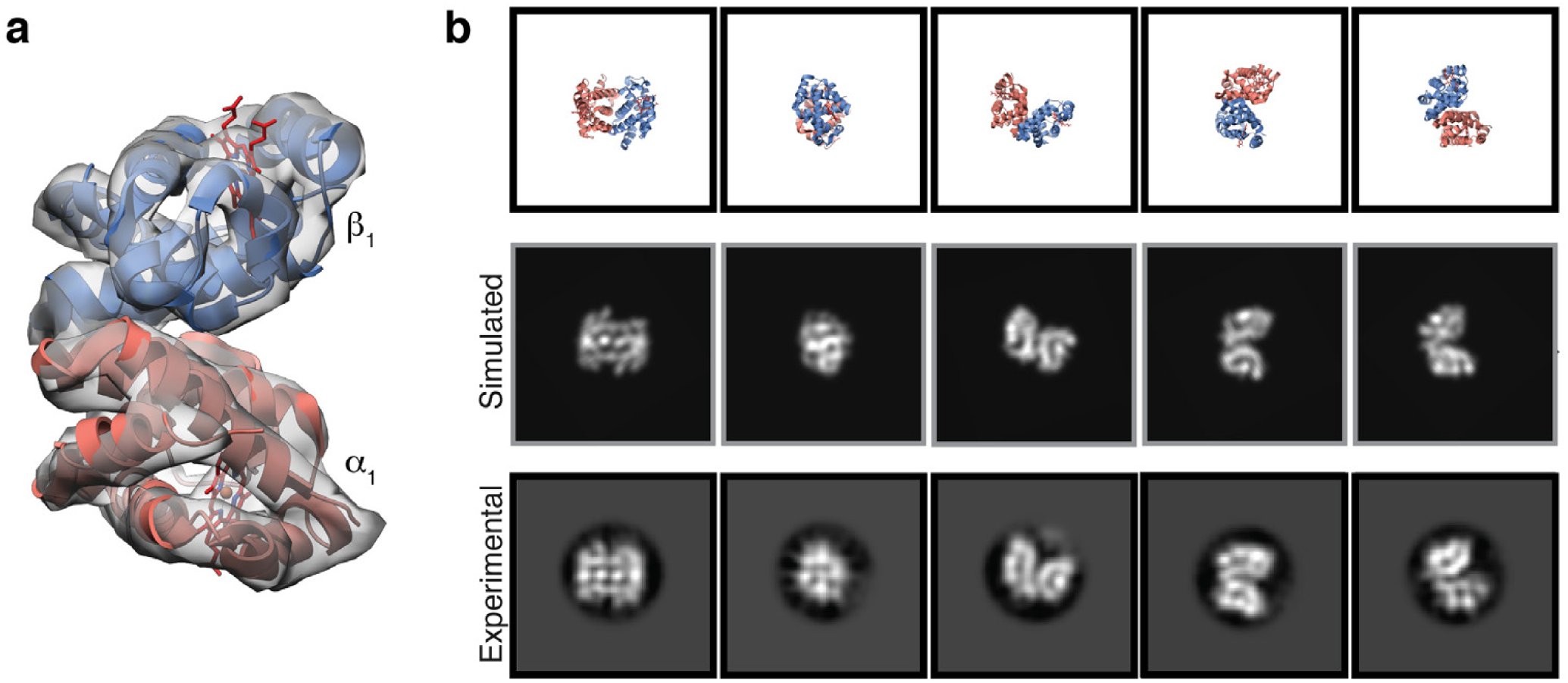
2D classification yields classes of ∼32 kDa hemoglobinαβ heterodimer. **(a)** Simulated EM density of the hemoglobin αβ heterodimer low-pass filtered to 8 Å resolution shown as a gray transparent surface. The αβ heterodimer atomic model is shown as ribbon cartoon and the hemes are shown as red sticks. **(b)** Views of the hemoglobin αβ heterodimer atomic model (top) and 2D projections of the simulated hemoglobin αβ heterodimer EM density (middle) corresponding to the class averages obtained from 2D classification (bottom) (see Figure 2).

We next decided to pursue the structure of the asymmetric ∼43 kDa catalytic domain of protein kinase A (PKAc) bound to ATP and IP20, an inhibitory pseudo-peptide substrate that has been previously shown to stabilize PKAc^25^. EM grid preparation and imaging of frozen-hydrated inhibited PKAc (iPKAc) were performed similarly as described for ADH and metHb with minor modifications (see Methods). Remarkably, given the small size and dimensions of iPKAc (65 *×* 40 *×* 40 Å), particles were clearly discernible in the aligned micrographs even at modest underfocus values (e.g. *<*1000 nm) (Fig. 5).

**Figure 5.**
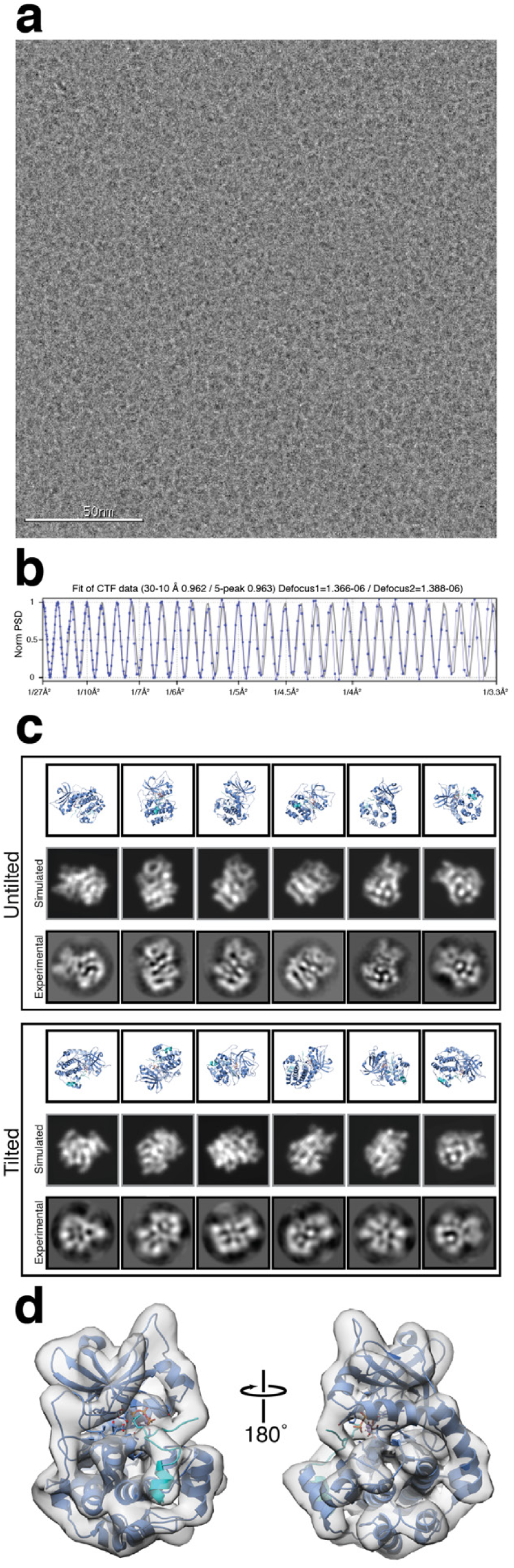
Towards a high-resolution cryo-EM reconstruction of the ∼43 kDa isolable kinase domain of protein kinase A. **(a)** Representative motion-corrected electron micrograph of the catalytic subunit of protein kinase A bound to IP20 (iPKAc) embedded in vitreous ice recorded at ∼1.3 *μ*m defocus (scale bar, 50 nm). **(b)** 1-dimensional plot of the contrast transfer function (CTF) Thon rings (black line) and the CTF estimated with CTFFIND4^33^ (blue line). **(c)** Views of the iPKAc atomic model (top, PDB ID: 1ATP, blue cartoon) and 2D projections of the simulated iPKAc EM density (middle) corresponding to the class averages obtained from 2D classification (bottom). **(d)** Final ∼6 Å iPKAc EM density shown as a transparent gray surface with the fitted atomic model (PDB ID: 1ATP) shown as a blue cartoon. ATP is shown as sticks and IP20 is colored light blue.

Reference-free 2D classification of gaussian-picked particles yielded featureful 2D class averages representing numerous views of iPKAc wherein the N- and C-terminal lobes of the complex, as well as secondary structural elements, could be clearly discerned. Consistent with these observations, ab initio 3D model generation using cryoSPARC^26^ yielded a volume resembling low-resolution iPKAc (Supplementary Fig. 3c). However, subsequent 3D auto-refinement yielded a *>*4 Å resolution reconstruction of iPKAc exhibiting pronounced features consistent with preferred orientation of particles at the air-water interface (e.g. stretching along the axis orthogonal to the dominant view) and over-fitting (e.g. streaking density extending beyond the masked region), indicating that the FSC-reported resolution was substantially inflated. Examination of the Euler distribution plot confirmed that regions of Euler space were unaccounted for (Supplementary Fig. 4c). In an effort to obtain these missing views we imaged the specimen at a tilt angle of 30^*°*^ using methodologies previously described^27^ (see Methods). 2D classification of particles extracted from the tilted data gave rise to averages comprising views that were not observed in the untilted data (Fig. 5c). However, the overall visual quality of the class averages from the tilted data were lower than those of the untilted data, presumably due to the combination of increased beam-induced motion and decreased SNR of the particles resulting from imaging a tilted specimen^27^. Nonetheless, our ability to obtain featureful iPKAc 2D class averages from tilted micrographs indicates that this imaging scheme can be applied to biological macro-molecules of a wide range of sizes.

3D auto-refinement of the combined particles from the tilted and untilted data yielded a ∼6 Å resolution reconstruction of sufficient quality to discern tertiary structure elements consistent with the iPKAc kinase fold. Although this reconstruction appeared to be more isotropic in resolution than the untilted data, the secondary structure elements of the EM density are smooth and featureless. Numerous computational efforts were employed (see Methods) to improve particle alignment and obtain a high-resolution reconstruction of iPKAc, but we were unable to overcome the resolution-limiting, gross misalignment of the particles (Supplementary Fig. 5). Given these complications arising from low image SNR, our work with iPKAc suggests that the use of a VPP and/or energy filter may potentially benefit imaging of targets of comparably small sizes (i.e. *<*50 kDa) by boosting observable contrast while preserving high spatial frequency information. However, the untilted 2D class averages demonstrate that detailed projections of *<*50 kDa particles are resolvable by conventional cryo-EM SPA, and that algorithmic development may enable a high-resolution structure to be produced from these data.

### Further developments

The VPP has previously demonstrated excellent utility in resolving the structures of some small biological specimens^5,11^, and consequently is now widely perceived as a necessity for resolving smaller targets. The results presented in this study define a new frontier for target sizes that can feasibly be resolved using cryo-EM without the need for a phase plate. Although the use of significant underfocus (e.g. *>*2000 nm) has been speculated to be required for imaging macromolecules *<*200 kDa by conventional cryo-EM SPA, it is clear from this work that even molecules as small as ∼43 kDa can be resolved with modest underfocus (e.g. *<*1000 nm) provided the vitreous ice encompassing the molecule of interest is sufficiently thin. Moreover, the results of our work with ADH and mHb demonstrate that small-molecule ligands and discrete conformational states within a single sample can be resolved using conventional SPA even for smaller complexes. Taken together, these findings broaden the potential of cryo-EM as a powerful tool for a variety of structure-based studies, particularly in drug discovery.

Though we could obtain a 3D reconstruction of iPKAc that allowed us to discern tertiary structural features, we were ultimately unable to achieve resolutions comparable to those of our ADH and mHb reconstructions. This was likely due to the lack of robust or accurate angular assignments of iPKAc particle images arising from the inherently limited SNR of the molecule in vitreous ice that was further confounded by molecule asymmetry, manifesting as gross misalignment within the 3D reconstruction. Future developments in detector technology and/or image processing algorithms will be integral for high-resolution structure determination of comparably small complexes. These improvements will also be of great relevance to structural studies of small membrane-embedded proteins, the SNRs for which are detrimentally impacted by the presence of disordered detergent micelles or nanodiscs. Our work further propels an important trajectory for cryo-EM SPA and indicates that high-resolution structure determination of complexes approaching, or even exceeding, the theoretical size limit of cryo-EM SPA will likely be realized in the near future.

## Methods

### Sampling Handling and Protein Purification

#### Alcohol Dehydrogenase

Lyophilized horse liver alcohol dehydrogenase (Sigma Aldrich) (ADH) was solubilized in 20 mM HEPES (pH 7.5), 10 mM NaCl, 1 mM TCEP (Buffer A) and dialyzed overnight against the same buffer at 4^*°*^ C. Dialyzed protein was then diluted two-fold with Buffer A lacking NaCl and immediately loaded onto a HiTrap SP HP (GE Life Sciences) that had been equilibrated in Buffer A. ADH was eluted using a gradient from 0% to 100% Buffer B (20 mM HEPES (pH 7.5), 500 mM NaCl, 1 mM TCEP) over 15 mL while collecting 500 *μ*L fractions. Fractions containing ADH were pooled and dialyzed overnight at 4^*°*^ C against 20 mM Tris (pH 8.5), 100 mM NaCl, 1 mM TCEP, and 0.5 mM NADH (Buffer C). ADH was then concentrated to ∼300 *μ*L using a 30,000 MWCO spin concentrator and subjected to size-exclusion chromatography using a Superdex 10/300 GL (GE Life Sciences) that had been equilibrated in Buffer C. Pure ADH was pooled and concentrated to ∼2.5 mg/mL and used immediately for cryo-EM grid preparation.

#### Hemoglobin

Lyophilized human methemoglobin (Sigma-Aldrich) (Hb) was solubilized in PBS (pH 7.5) to a final concentration of ∼12 mg/mL and used without further modifications. A UVVIS absorbance spectrum of the solubilized protein indicated a Soret maximum at 406 nm, consistent with previous measurements of methemoglobin, but with two small peaks in the α/β region at 576 nm and 541 nm, respectively, potentially indicating a small amount of deoxyhemoglobin in the sample.

#### Protein Kinase A

The catalytic subunit of PKA was expressed, purified, and bound to ATP and IP20 according to protocols described previously^28^. iPKAc was donated by P. Aoto of the Taylor lab (UCSD) and used without modification.

### Cryo-EM Grid Preparation

3 *μ*L of purified alcohol dehydrogenase (∼2.5 mg/mL, methemoglobin (∼12 mg/mL, or inhibited PKA (∼5 mg/mL) were dispensed on UltrAuFoil R1.2/1.3 300-mesh grids (Electron Microscopy Services) that had been freshly plasm cleaned using a Solarus plasma cleaner (Gatan, Inc.) with a 75% argon / 25% oxygen atmosphere at 15 Watts for 6 seconds. Grids were immediately blotted manually^29^ using a custom-built manual plunger in a cold room (≥95% relative humidity, 4^*°*^ C)^12^. Samples were blotted for 4-5 s with Whatman No. 1 filter paper immediately before plunge freezing in liquid ethane cooled by liquid nitrogen. We aimed to achieve a particle concentration that maximized the number of particles contained within the holes without resulting in overlapping particles and/or aggregation in order to maintain a consistent ice thickness across the center of holes and provide sufficient signal for accurate CTF estimation, as observed previously^12^.

### Cryo-EM Data Acquisition, Image Processing, and Refinement

Microscope alignments were performed on a cross-grating calibration grid using methodologies previously described^12,14^ with minor modifications. Briefly, after obtaining parallel illumination in diffraction mode at a diffraction length of D 850 mm, the length was increased to D 5.7 m and the intensity was adjusted to minimize the size of the caustic spot followed by minimization of the astigmatism of the diffraction lens. This process was iterated until no further improvements could be discerned visually. The resulting beam-intensity value was saved in Leginon for the exposure preset (73000x, 0.56 Å /pixel) and remained unchanged throughout data collection. The objective aperture was then centered, objective lens astigmatism was minimized, and coma-free alignment was performed using Leginon^13^ as described previously^12,30^. Daily adjustments were made, if necessary, during data collection to maintain lens stigmation and to ensure the beam was centered properly. The hardware darks of the K2 Summit DED were updated approximately every 8-12 hours.

All cryo-EM data were collected on a Thermo Fisher Talos Arctica transmission electron microscope (TEM) operating at 200 keV. All cryo-EM data were acquired using the Leginon^13^ automated data-collection program and all image pre-processing (e.g. frame alignment, CTF estimation, and initial particle-picking) were performed in real-time using the Appion^31^ image-processing pipeline. Movies were collected using a K2 Summit DED (Gatan, Inc.) operating in counting mode (0.56 Å /pixel) at a nominal magnification of 73,000x using a defocus range of −0.5 *μ*m to −1.6*μ*m. Movies were collected over an 11 second exposure with an exposure rate of ∼1.95 e^-^/pixel/s, resulting in a total exposure of ∼69 e^-^ Å ^-2^ (1.57 e^-^ Å ^-2^/frame). Motion correction and dose-weighting were performed using the MotionCor2 frame alignment program^32^ as part of the Appion pre-processing workflow. Frame alignment was performed on 5 *×* 5 tiled frames with a B-factor of 100 (ADH and Hb) or 250 (PKA) applied. A running average of 3 frames was also used for PKA for frame alignment.

Unweighted summed images were used for CTF determination using CTFFIND4^33^ and gCTF^34^. Specifically, CTFFIND4 within Appion was used for real-time CTF determination (512 box size, 40 Å minimum resolution, 3 Å max resolution, 0.10 amplitude contrast), wherein aligned micrographs with a CTF estimate confidence of fit below 90% were eliminated from further processing. Local CTF estimates using gCTF were then performed on the remaining micrographs using using the standalone package (ADH) or a grid-based algorithm incorporated within Appion (metHb, PKA). For the latter, dummy coordinates were placed across the micrograph in rows and columns spaced 200 pixels apart and passed to gCTF for local CTF estimation using equiphase averaging (local box size of 512 pixels, 0.10 amplitude contrast, 50 Å minimum resolution, 4 Å max resolution, 1024-pixel field size). CTF values for a given particle was then determined using a cubic spline interpolation of the local CTF estimates within the grid.

Local resolution estimates for all reconstructions were calculated using the blocres function in BSOFT^23^.

#### Alcohol Dehydrogenase

Difference of Gaussian (DoG) picker^35^ was used to automatically pick particles from the first 145 dose-weighted micrographs yielding a stack of 135,424 picks that were binned 4 *×* 4 (2.23 Å /pixel, 64 pixel box size) and subjected to reference-free 2D classification using RELION 2.1^19^. The best five classes that represented orthogonal views of ADH were then used for template-based particle picking using RELION. 1,232,543 picks were extracted from 1,151 dose-weighted movies, binned 4 *×* 4 (2.23 Å /pixel, 64-pixel box size) and subjected to reference-free 2D classification using RELION using a 110 Å soft circular mask. The best 2D class averages that represented side or top/bottom views of ADH (i.e. the longest dimensions of ADH, see Figure 1) were then isolated (639,430 particles). Those class averages that contained “end-on” or tilted views of ADH (e.g. the smallest dimension of ADH, see Figure 1) were combined and subjected to another round of reference-free 2D classification using a 60 Å soft circular mask. The best 2D class averages were then selected (20,232 particles) were then combined with the long views for further processing.

A total of 659,662 particles corresponding to the best 2D class averages that displayed strong secondary-structural elements and multiple views of ADH were selected for homogenous ab inito model generation using cryoSPARC^26^ to eliminate potential model bias. The generated model exhibited C2 symmetry and was low-pass filtered to 30 Å for use as an initial model for 3D auto-refinement in RELION. 659,662 binned 4 *×* 4 particles (2.23 Å /pixel, 64-pixel box size) were 3D auto-refined into a single class followed by subsequent re-centering and re-extraction binned 2 *×* 2 (1.12 Å /pixel, 256-pixel box size). Due to the close proximity of neighboring particles, any re-centered particle within a 30-pixel range of another was considered a duplicate and subsequently removed. These particles were then 3D auto-refined (C2 symmetry) into a single class using a scaled version of the binned 4 *×* 4 refined map. Upon convergence, the run was continued with a soft mask (5-pixel extension, 5-pixel soft cosine edge) followed by a no-alignment, 3D classification (6 classes, tau fudge=20) using the same soft mask. Particles comprising the best-resolved class was then subjected to 3D auto-refinement (C2 symmetry) using a soft mask. A subsequent no-alignment classification (2 classes, tau fudge=20) was performed and the class that possessed the best-resolved side-chain and backbone densities were re-centered and re-extracted unbinned (0.56 Å /pixel, 512-pixel box size). This final stack of 11,672 particles was 3D auto-refined using a soft mask to a final estimated resolution of 2.92 Å (C2 symmetry) and 3.45 Å (C1 symmetry) (gold-standard FSC at 0.143 cutoff) using phase randomization to account for the convolution effects of a solvent mask on the FSC between the two independently refined half maps^22,36^.

#### Methemoglobin

DoG picker^35^ was used to automatically pick particles from the first 560 dose-weighted micrographs yielding a stack of 234,895 picks that were binned 4 *×* 4 (2.23 Å /pixel, 64 pixel box size) and subjected to reference-free 2D classification using RELION^19^. The best nine classes that represented orthogonal views of Hb were then used for template-based particle picking using RELION. 1,615,738 picks were extracted from 1,673 dose-weighted movies, binned 4 *×* 4 (2.23 Å /pixel, 64-pixel box size) and subjected to reference-free 2D classification using RELION using a 80 Å soft circular mask. Class averages that displayed strong secondary-structural elements of metHb (513,632 particles) were combined and subjected to another round of reference-free 2D classification. A total of 160,169 particles corresponding to the best 2D class averages that displayed multiple views of tetrameric Hb were selected for homogenous ab inito model generation using cryoSPARC^26^ to eliminate potential model bias. The generated volume was low-pass filtered to 20 Å and used as an initial model for 3D auto-refinement in RELION.

Binned 4 *×* 4 particles (2.23 Å /pixel, 64-pixel box size) were 3D auto-refined into a single class followed by subsequent re-centering and re-extraction binned 2 *×* 2 (1.12 Å /pixel, 128-pixel box size). Any re-centered particle within a 25-pixel range of another was considered a duplicate and subsequently removed. These particles were then 3D auto-refined (C2 symmetry) into a single class using a scaled version of the binned 4 *×* 4 refined map. Upon convergence, the run was continued with a soft mask (5-pixel extension, 5-pixel soft cosine edge) followed by three parallel no-alignment 3D classifications (four classes) with varying tau fudge values (8, 12, or 20) using the same soft mask. Particles comprising the best-resolving classes across each classification were combined (35,809 particles) and duplicates were eliminated. The particles were then subjected to no-alignment classification (2 classes, tau fudge=20). Each class was then separately refined, re-centered, and re-extracted unbinned (0.56 Å /pixel, 256-pixel box size) and 3D auto-refined using a soft mask. Class 1 (24,308 particles) refined to a final estimated resolution of ∼2.8 Å (C2 symmetry, ∼3.2 Å resolution with C1 symmetry) and Class 2 (11,501 particles) refined to a final estimated resolution of ∼3.2 Å resolution (C2 symmetry, ∼3.5 Å resolution with C1 symmetry) according to gold-standard FSC^21^ using phase randomization to account for the convolution effects of a solvent mask on the FSC between the two independently refined half maps^22,36^.

#### Protein Kinase A

DoG picker^35^ was used to automatically pick particles from the untilted, aligned and dose-weighted micrographs yielding a stack of 554,170 picks that were binned 4 *×* 4 (2.23 Å /pixel, 64-pixel box size) and subjected to reference-free 2D classification using RELION^19^. A total of 314,001 particles corresponding to the best 2D class averages that displayed multiple views of iPKAc were selected for homogenous ab inito model generation using cryoSPARC^26^ to eliminate potential model bias. The generated volume was low-pass filtered to 20 Å and used as an initial model for 3D auto-refinement (C1 symmetry) in RELION. Particles were then re-centered and re-extracted, Fourier binned 2 *×* 2 (1.12 Å /pixel, 128-pixel box) and subsequently 3D classified (tau fudge=4, E-step limit=7 Å). Particles comprising the best-resolved class was 3D auto-refined to ∼4.3 Å resolution, as estimated by goldstandard FSC. However, this EM density exhibited artifacts associated with preferred orientation (stretched features in directions orthogonal to the preferred view) and did not possess high-resolution features consistent with the Gold-standard FSC-estimated resolution of ∼4.3 Å resolution. In an attempt to obtain the missing views of iPKAc and potentially improve the resolution of our reconstruction we collected a dataset at a tilt angle of 30^*°*^ using methodologies previously described^27^.

These aligned, dose-weighted micrographs were combined with the untilted data and forward projections of a 10 Å low-pass filtered EM density of iPKAc were used for template-based particle picking within RELION. Together, 1,762,088 particles were extracted, Fourier binned 4 *×* 4 (2.23 Å /pixel, 64-pixel box), and subjected to reference-free 2D classification (tau fudge=1, E-step limit=7 Å) using RELION. It was apparent from the resulting 2D class averages that additional views of iPKAc were obtained from imaging a tilted specimen (Figure 5). Particles comprising the “best” classes were 3D classified (tau fudge=1, E-step limit=7 Å) and those classes that most resembled iPKAc were subjected to an additional round of 3D classification (tau fudge=1, E-step limit=7 Å). The “best” resolved classes were then 3D auto-refined (C1 symmetry) to yield a ∼4.6 Å resolution reconstruction, as estimated by gold-standard FSC. Although these particles refined to a similar resolution as the untitled data alone, the resulting reconstruction was more isotropic. Particles were re-centered and re-extracted, Fourier binned 2 *×* 2 (1.12 Å /pixel, 128-pixel box) and duplicate particle picks were eliminated. These particles were 3D auto-refined to ∼6.2 Å resolution and subjected to no-alignment 3D classification (tau fudge=6). The “best” resolved classes were combined and 3D-auto-refined to ∼6.2 Å resolution. Refined particles with NrOfSignificantSamples values *>*25 were eliminated and the remaining particles were 3D auto-refined to ∼ 4.3 Å resolution (as estimated by gold-standard FSC).

Although the additional views from the tilted dataset aided in lessening the artifacts resulting from preferred orientation, the final EM density does not possess molecular features consistent with the reported resolution of ∼4.3 Å but rather resembles a 6-7 Å resolution EM density. Strategies to further improve the quality of the reconstruction did not yield significant results. Such attempts included: 1) high-pass filtering the extracted particles to 100 or 120 Å prior to 3D refinements as has previously been implemented for high-resolution structure determination of small proteins^11^; 2) increasing the tau fudge value (e.g. 4, 6, 8, etc.) during refinement to place greater weight on the experimental data; 3) increasing the amplitude contrast of the inputted particle stack (e.g 0.2 vs. 0.10 as used for the final structure); 4) filtering the particles iteratively by RELION metadata metrics, e.g. NrOfSignificantSamples, MaxValueProbDistribution, or Z-score; 5) using particle images with a box size just outside the bounds of the particle (e.g. 72 pixel box vs. 128 pixel box for binned 2 *×* 2 particles); or 6) various combinations of the above mentioned.

### Model Building and Refinement

For each of the final reconstructions, an initial model that had been stripped of all ligands, waters, and alternative conformations, had all occupancies set to zero, had a single B-factor value was set for all atoms, and had all Ramachandran, rotameric, and geometric outliers corrected, was subjected to a multi-model pipeline using methodologies similar to those previously described^37^. Briefly, PDB ID: 2JHF was used as the starting model for ADH, chains A-D from PDB ID: 4N7P was used for Hb class 1, and chains E-H from PDB ID: 4N7N were used for Hb class 2. These initial models were then refined into the EM density using Rosetta^38^ while enforcing the same symmetry that was applied during reconstruction and adjusting the Rosetta weighting and scoring functions according to the FSC-estimated map resolution. Each of the 200 Rosetta-refined models were then ranked based on the number of Ramachandran outliers, geometry violations, Rosetta aggregate score, and MolProbity clashscore^39^. The 10 structures that scored the best across all categories were selected for further real-space refinement using the Phenix refinement package^40^ after incorporating cofactors (e.g. NADH for ADH and heme for Hb) and active-site and structural Zn ions (ADH). Model–model agreement statistics were calculated using a previously described approach^37^.

## Acknowledgements

We thank J.C. Ducom at The Scripps Research Institute (TSRI) High Performance Computing facility for computational support, B. Anderson at the TSRI electron microscopy facility for microscope support, and M. Vos for advice and discussion regarding TEM alignments. We thank P. Aoto of the S. Taylor laboratory (University of California, San Diego) for kindly providing iPKAc for this study. G.C.L. is supported as a Searle Scholar and as a Pew Scholar, by a young investigator award from Amgen, and by the US National Institutes of Health (NIH) grant DP2EB020402. Computational analyses of EM data were performed using shared instrumentation at TSRI funded by NIH S10OD021634. M.A.H. is supported by a Helen Hay Whitney Foundation postdoctoral fellowship. M.W is supported by a National Science Foundation Graduate Student Research Fellowship.

## Author Contributions

M.A.H. and M.W. performed all cryo-EM experiments and analyses under the supervision of G.C.L. All authors contributed to the experimental design and manuscript preparation.

## Data Availability

The atomic coordinates for the ADH and Hb (class 1 and class 2) structures have been deposited in the Protein Data Bank (PDB) under accession codes 6NBB, 6NBC, and 6NBD, respectively. The corresponding EM density maps (final unsharpened and sharpened maps, half maps, and masks) have been deposited to the Electron Microscopy Data Bank under accessions EMD-0406, EMD-0407, and EMD-0408, respectively, and EMD-0409 for iPKAc. Uncorrected movie frames and associated gain correction images for all the datasets in this study are currently being uploaded to the Electron Microscopy Public Image Archive (EMPIAR) and will be made available.

## Supplementary Materials

**Supplementary Figure 1.**
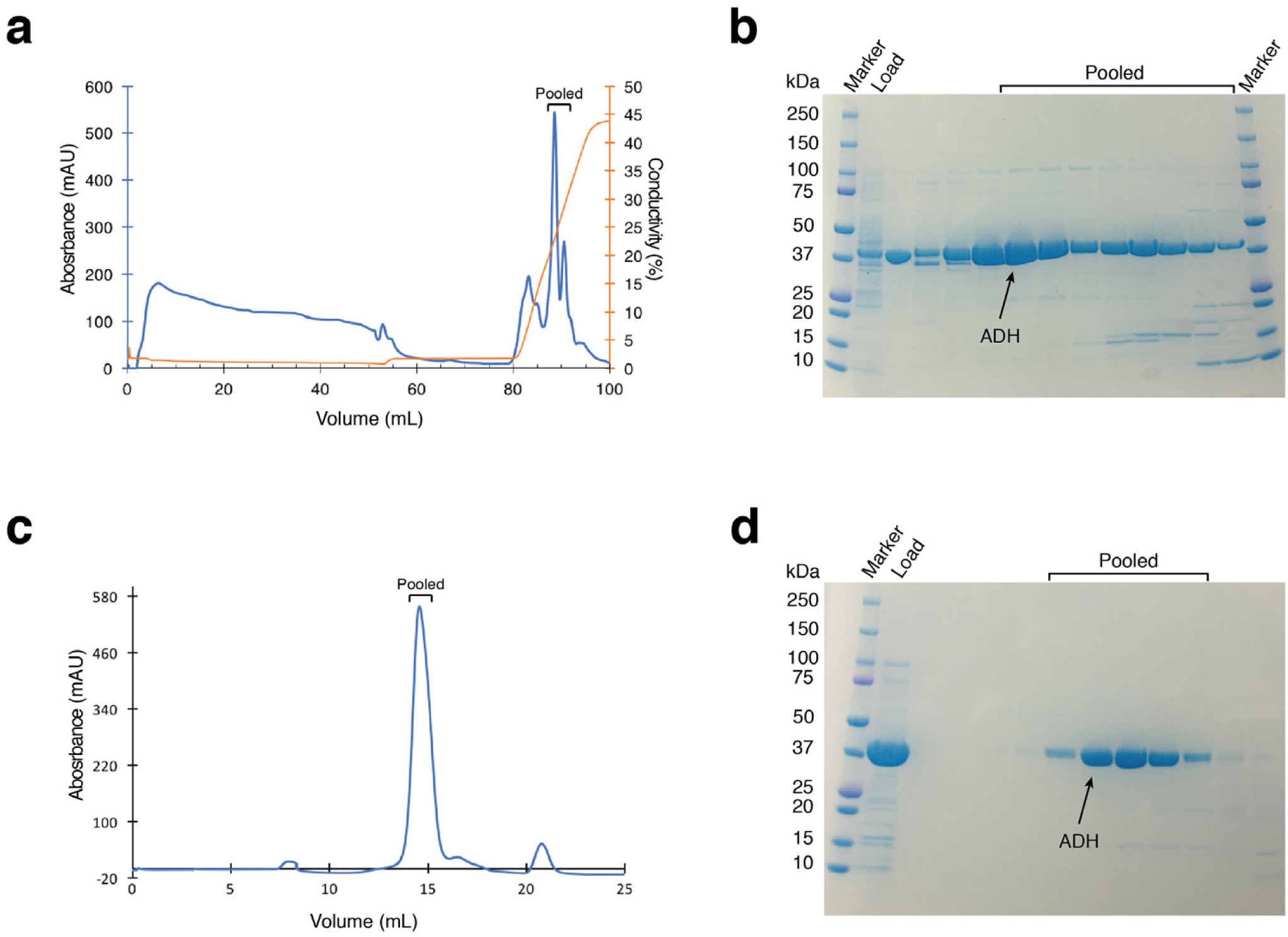
Purification of alcohol dehydrogenase. **(a)** Elution profile of alcohol dehydrogenase (ADH) from a HiTrap SP sepharose cation exchange column. Traces for UV absorbance (mAU) (blue) and conductivity (%) (orange) are shown. Fractions that were pooled are indicated. **(b)** SDS-PAGE analysis of ADH elution from HiTrap SP sepharose shown in (a). Molecular weight marker, load, and pooled fractions are indicated. **(c)** Elution profile of ADH from a Superdex 200 gel filtration column. UV absorbance (mAU) is shown as a blue trace. Fractions that were pooled are indicated. **(d)** SDS-PAGE analysis of the gel filtration run shown in (c). Molecular weight marker, load, and pooled fractions are indicated.

**Supplementary Figure 2.**
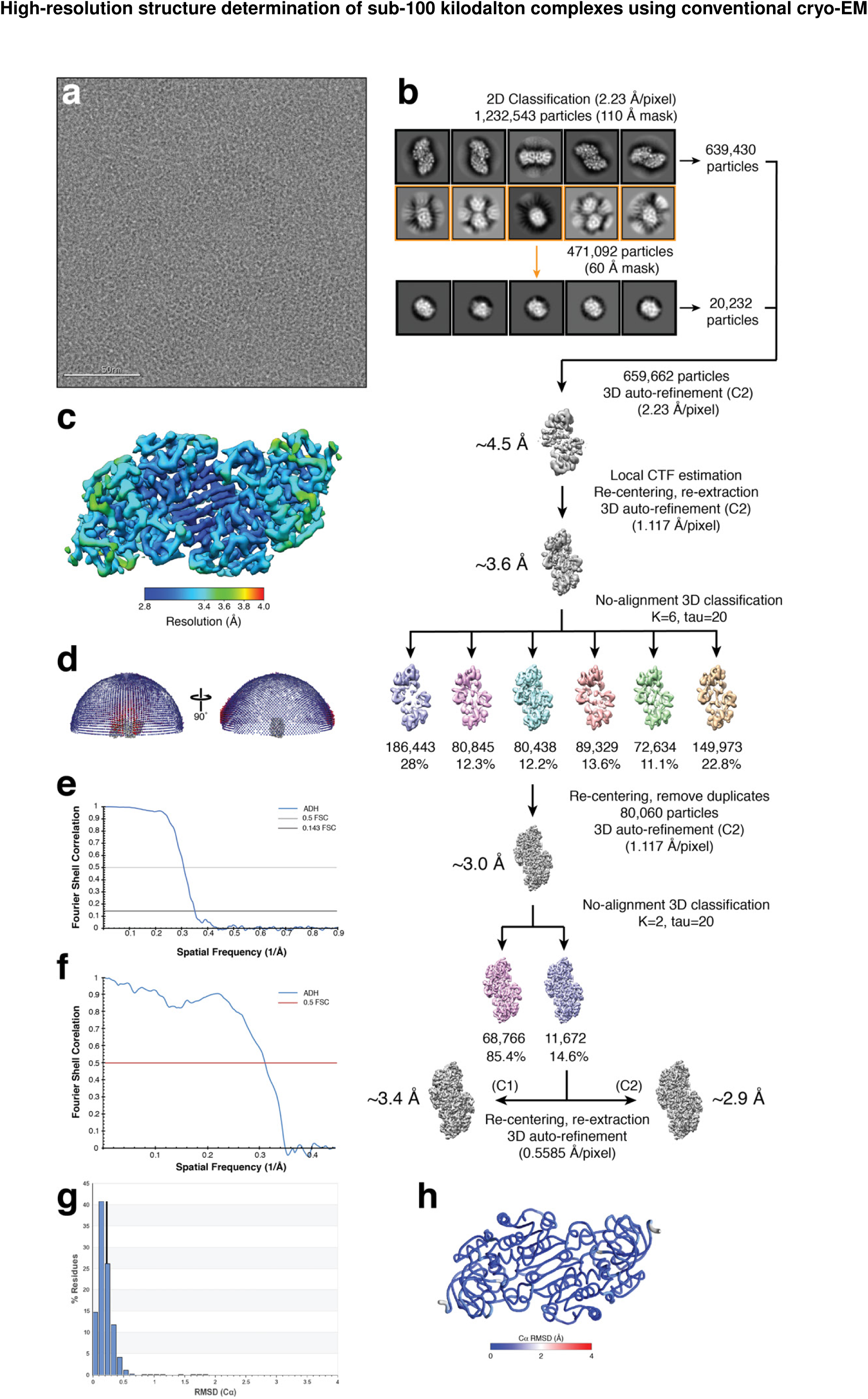
Schematic for alcohol dehydrogenase single-particle cryo-EM data processing. **(a)** Representative motioncorrected micrograph of vitrified ADH collected at ∼ 1 *μ*m underfocus. **(b)** ∼ 1.2 million particles were extracted from the aligned, dose-weighted micrographs, Fourier binned 4 *×* 4, and subjected to two subsequent rounds of reference-free 2D classification using RELION^18^. 2D classes highlighted in yellow were selected for a second round of 2D classification using a smaller soft circular mask (60 Å vs. 110 Å) to discern orthogonal views of ADH. Representative 2D class averages are shown. Particles comprising the “best” classes were 3D auto-refined to ∼4.5 Å resolution followed by local CTF estimation, particle coordinate re-centering, and re-extraction Fourier binned 2 *×* 2. 3D auto-refinement of these particles yielded a ∼3.6 Å resolution reconstruction that was then subjected to no-alignment 3D classification (tau fudge=20) and the best resolved class (80,060 particles) was further auto-refined to ∼3.0 Å resolution. Following another no-alignment 3D classification, particles (11,672) composing the “best” resolved class were re-centered and re-extracted unbinned, and auto-refined to ∼2.9 Å resolution (C2 symmetry) or ∼3.4 Å resolution (C1 symmetry). **(c)** ADH EM density colored by local resolution (estimated using BSOFT^23^). **(d)** Plot showing the Euler angle distribution of the final ADH EM density. **(e)** Gold-standard FSC curve generated from the independent half maps contributing to the ∼2.9 Å resolution ADH EM density. **(f)** FSC curve calculated between the ADH EM density and the refined atomic model. **(g)** Histogram of the per-residue Cα RMSD values calculated from the top 10 refined ADH atomic models. **(h)** Worm plot representation of the per-residues Cα RMSD calculated from the top 10 refined ADH atomic models.

**Supplementary Figure 3.**
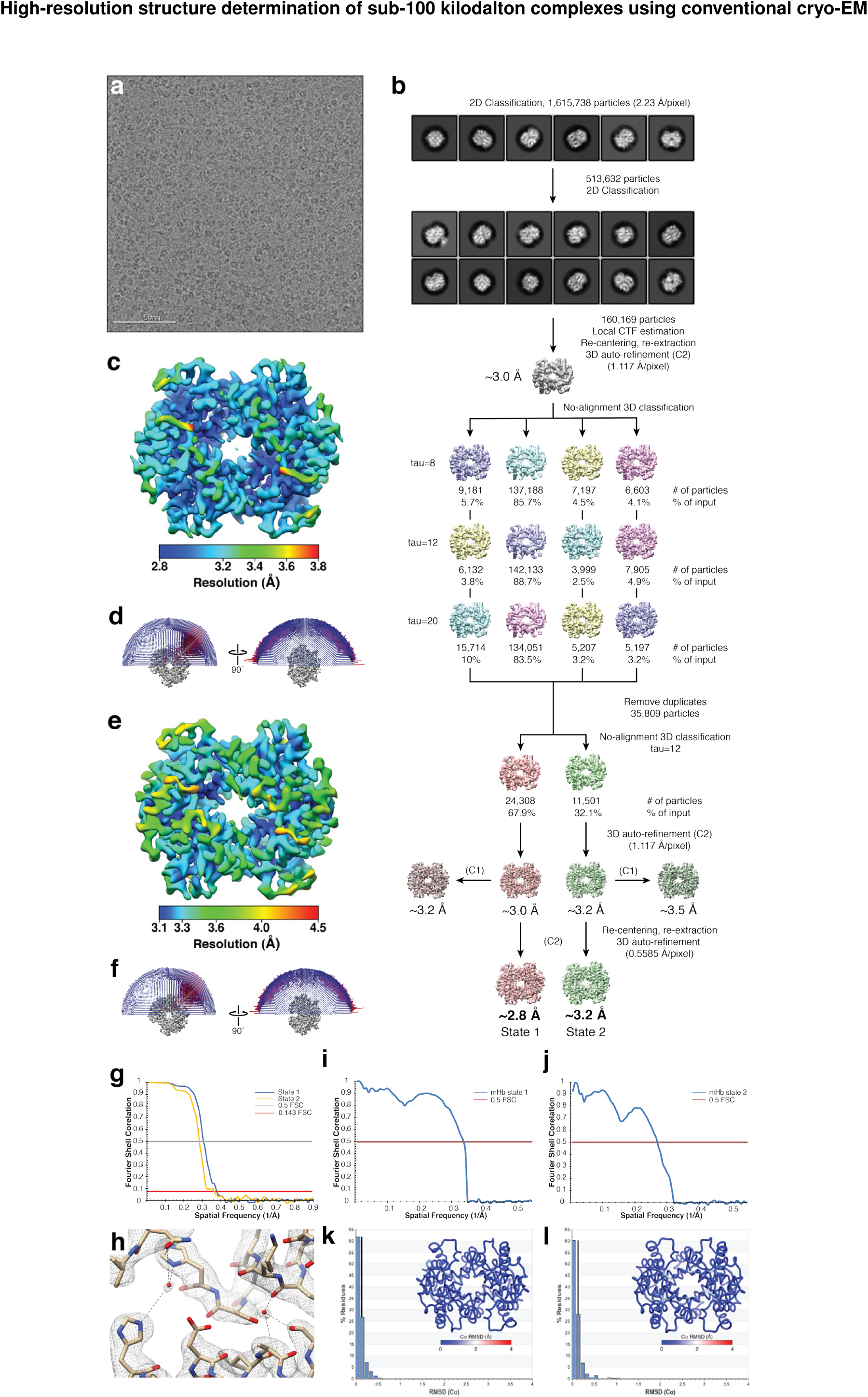
Schematic for methemoglobin single-particle cryo-EM data processing. **(a)** Representative motion-corrected micrograph of vitrified methemoglobin (mHb) collected at ∼1 *μ*m underfocus. **(b)** ∼1.6 million particles were extracted from the aligned, dose-weighted micrographs, Fourier binned 4 *×* 4, and subjected to two subsequent rounds of reference-free 2D classification using RELION^18^. Representative 2D class averages are shown. Particles comprising the “best” classes (160,169) were used for local CTF estimation, particle coordinate re-centering, and re-extraction Fourier binned 2 *×* 2. 3D auto-refinement of these particles yielded a ∼3.0 Å resolution reconstruction that was then subjected to three parallel no-alignment 3D classifications (tau fudge values of 8, 12, or 20). Unique particles corresponding to the best resolved classes from each classification were combined (35,809 particles) and further 3D auto-refined to ∼3.0 Å resolution. Another no-alignment 3D classification of these particles yielded two conformationally distinct classes (state 1 and state 2) that 3D auto-refined to ∼2.8 and ∼3.2 Å resolution, respectively, with C2 symmetry applied. **(c)** and **(e)** mHb state 1 and state 2 EM densities, respectively, colored by local resolution (estimated using BSOFT^23^). **(d)** and **(f)** Plots showing the Euler angle distribution for the final mHb state 1 and state 2 EM densities, respectively. **(g)** Gold-standard FSC curves generated from the independent half maps contributing to the ∼2.8 Å (state 1, blue trace) or the ∼3.2 Å resolution (state 2, orange trace) mHb EM densities **(h)** Ordered water molecules (red spheres) in the mHb state 1 EM density (gray mesh). Putative hydrogen bonds to the water molecules are shown as black dotted lines. **(i)** FSC curve calculated between the mHb state1 EM density and the refined atomic model. **(j)** FSC curve calculated between the mHb state 2 EM density and the refined atomic model. **(k)** Histogram of the per-residue Cα RMSD values calculated from the mHb state 1 top 10 refined atomic models. Inset is a worm plot representation of per-residue Cα RMSD calculated from the top 10 refined mHb state 1 atomic models. **(l)** Histogram of the per-residue Cα RMSD values calculated from the mHb state 2 top 10 refined atomic models. A worm plot representation of per-residue Cα RMSD calculated from the top 10 refined mHb state 2 atomic models is shown (inset).

**Supplementary Figure 4.**
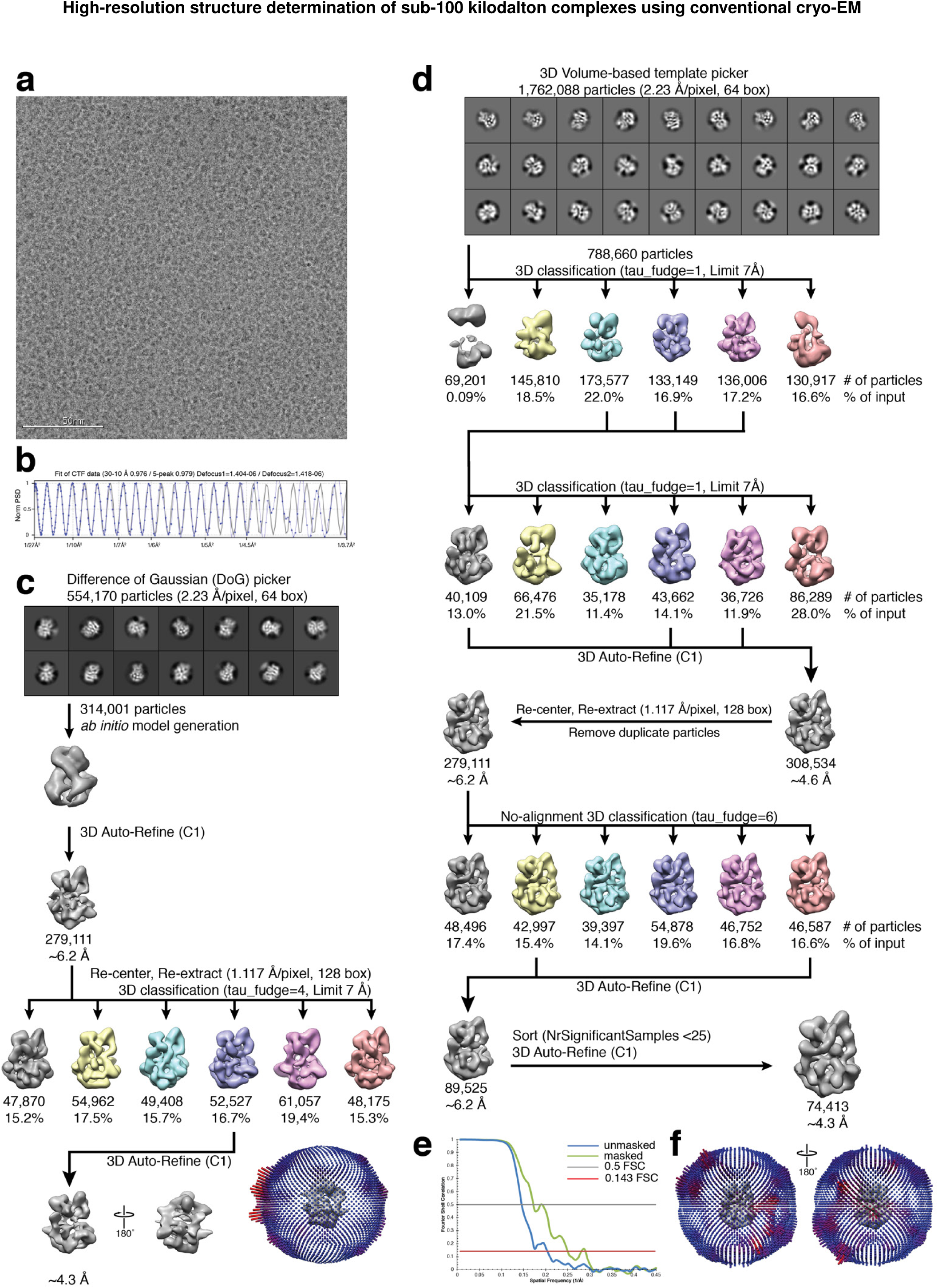
Schematic for protein kinase A catalytic domain-IP20 complex single-particle cryo-EM data processing. **(a)** Representative motion-corrected micrograph of vitrified iPKAc tilted at 30^*°*^ and imaged at ∼1.4 *μ*m underfocus. **(b)** 1-dimensional plot of the contrast transfer function (CTF) Thon rings (black line) and the CTF estimated with CTFFIND4^33^ (blue line). **(c)** ∼554K DoG-picked particles obtained from untilted data collection were extracted from the aligned, dose-weighted micrographs, Fourier binned 4 *×* 4 (2.23 Å /pixel, 64-pixel box), and subjected to reference-free 2D classification using RELION^18^. Representative 2D class averages are shown. Particles comprising the “best” classes were 3D auto-refined (C1 symmetry) using an ab initio model created using cryoSPARC^26^. Particles were re-centered and re-extracted, Fourier binned 2 *×* 2 (1.12 Å /pixel, 128-pixel box) and subsequently 3D classified (tau fudge=4, E-step limit=7 Å). The best-resolved class was 3D auto-refined to ∼4.3 Å resolution, as estimated by gold-standard FSC. Plot showing the Euler angle distribution. **(d)** ∼1.76 million particles from the combined untilted and tilted data, picked using a 3D-volume-based template picker in RELION^18^, were extracted from the aligned and dose-weighted micrographs, Fourier binned 4 *×* 4 (2.23 Å /pixel, 64-pixel box), and subjected to RELION^18^ reference-free 2D classification. Representative 2D class averages are shown. Particles comprising the “best” classes were subjected to two rounds of 3D classification (tau fudge=1, E-step limit=7 Å). The “best” resolved classes were 3D auto-refined (C1 symmetry) to yield a ∼4.6 Å resolution reconstruction, as estimated by gold-standard FSC. Particles were re-centered and re-extracted, Fourier binned 2 *×* 2 (1.12 Å /pixel, 128-pixel box) and duplicate particle picks were eliminated. These particles were 3D auto-refined to ∼6.2 Å resolution and subjected to no-alignment 3D classification (tau fudge=6). The “best” resolved classes were combined and 3D-auto-refined to ∼6.2 Å resolution. Particles with NrOfSignificantSamples *>*25 were eliminated and 3D auto-refined to ∼4.3 Å resolution. **(e)** Gold-standard FSC curve generated from the independent half maps contributing to the ∼4.3 Å resolution iPKAc EM density. The “bumpiness” of the FSC is consistent with over-alignment of particle data, and the reported FSC resolution is not consistent with the quality of the reconstruction, which appears to be at ∼6 Å based on visual inspection. **(f)** Plots showing the Euler angle distribution of the final iPKAc EM density.

**Supplementary Figure 5.**
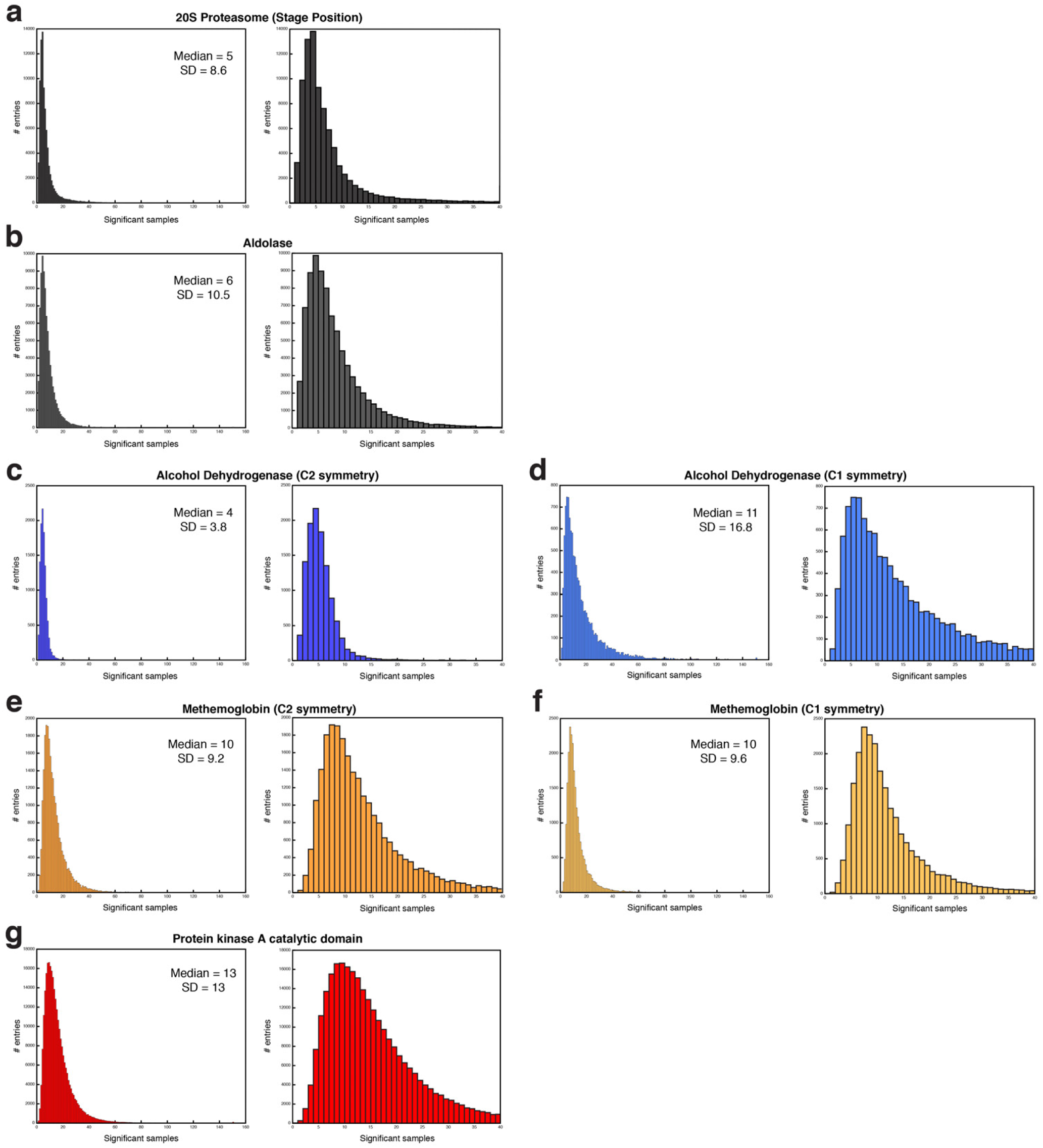
Distribution of RELION significant samples. Histogram plots of the number of significant samples of each particle contributing to the final cryo-EM reconstructions (left, 160 maximum; right, zoomed 40 maximum) of **(a)** 20S proteasome (EMDB ID: 8741), **(b)** aldolase (EMDB ID: 8743), **(c)** ADH C2-symmetric and **(d)** C1-symmetric refinements, **(e)** mHb State 1 C2-symmetric and **(f)** C1-symmetric refinements, and **(g)** iPKAc. The median and standard deviation values are reported.

**Supplementary Table 1.** Cryo-EM data collection, refinement, and validation statistics

|  | Alcohol Dehydrogenase | Hemoglobin |  | Protein Kinase A |
| --- | --- | --- | --- | --- |
| PDB ID | 6NBB | 6NBC | 6NBD | - |
| EMDB ID | EMD-0406 | EMD-0407 | EMD-0408 | EMD-0409 |
| **Data collection** |  |  |  |  |
| Microscope | Talos Arctica | Talos Arctica |  | Talos Arctica |
| Camera | K2 Summit | K2 Summit |  | K2 Summit |
| Magnification | 73,000 | 73,000 |  | 73,000 |
| Voltage (kV) | 200 | 200 |  | 200 |
| Exposure navigation | Stage Position | Stage Position |  | Stage Position |
| Cumulative exposure (e<sup>-</sup>/Å<sup>2</sup>) | 69 | 69 |  | 69 |
| Exposure rate (e<sup>-</sup>/pixel/sec) | 1.95 | 1.95 |  | 1.95 |
| Pixel size (Å/pixel) | 0.56 | 0.56 |  | 0.56 |
| Defocus range (μm) | 0.5 to 1.6 | 0.5 to 1.6 |  | 0.5 to 1.6 |
| Stage Tilt (°) | 0 | 0 |  | 0 30 |
| Micrographs collected (no.) | 1,151 | 1,673 |  | 3,176 2,402 |
| Total extracted picks (no.) | 1,232,543 | 1,615,738 |  | 667,212 1,504,429 |
| Refined particles (no.) | 659,662 | 160,169 |  | 1,760,088 |
| **Reconstruction** |  |  |  |  |
| Final Particles (no.) | 11,672 | 24,308 | 11,501 | 74,413 |
| Symmetry | C2 | C2 | C2 | C1 |
| Map sharpening B-factor (Å<sup>2</sup>) | -67 | -80 | -93 | -158 |
| Resolution (global) (Å) |  |  |  |  |
| FSC 0.5 (unmasked/masked) | 7.94/3.25 | 3.86/3.25 | 4.27/3.53 | 6.80/5.11<sup>†</sup> |
| FSC 0.143 (unmasked/masked) | 3.99/2.92 | 3.32/2.8 | 3.62/3.25 | 4.93/4.26<sup>†</sup> |
| Local resolution range (Å) | 2.8 - 13.9 | 2.6 - 10.4 | 3.2 - 10.1 | N/A |
| **Model composition** |  |  |  |  |
| Protein residues (atoms) | 768 (5570) | 566 (4472) | 566 (4472) |  |
| Ligands (atoms) | 6 (92) | 4 (176) | 4 (176) |  |
| **Reconstruction\*** |  |  |  |  |
| Map Correlation Coefficient |  |  |  |  |
| Global | 0.37 (0.37) | 0.61 (0.61) | 0.58 (0.58) |  |
| Local | 0.84 (0.83) | 0.85 (0.84) | 0.83 (0.81) |  |
| R.M.S. deviations |  |  |  |  |
| Bond lengths (Å) | 0.009 (0.010) | 0.009 (0.09) | 0.008 (0.009) |  |
| Bond angles (°) | 0.921 (0.951) | 1.003 (0.992) | 0.960 (0.979) |  |
| **Validation** |  |  |  |  |
| MolProbity score | 1.63 (1.60) | 1.03 (1.16) | 1.27 (1.32) |  |
| Clashscore (all atoms) | 5.17 (6.57) | 2.14 (2.69) | 3.71 (4.62) |  |
| Poor rotamers (%) | 0 (0.15) | 0.8 (0.44) | 0 (0.39) |  |
| Ramachandran |  |  |  |  |
| Favored (%) | 95.7 (95.5) | 96.4 (97.5) | 97.5 (97.5) |  |
| Allowed (%) | 4.3 (4.5) | 3.5 (2.5) | 2.5 (2.5) |  |
| Disallowed (Å) | 0 (0) | 0 (0) | 0 (0) |  |
| Mean per-residue Cα RMSD (Å) | 0.31 | 0.16 | 0.21 |  |
| Per-residue Cα RMSD range (Å) | 0.64 | 0.47 | 0.62 |  |
| EMRinger<sup>41</sup> Score | 3.90 (3.87) | 5.04 (5.12) | 4.87 (4.82) |  |

\* Values in parentheses correspond to values from top 10 models
<sup>†</sup> Reported values are not consistent with quality of density, likely inflated

## References

1. Cheng, Y. Single-particle cryo-EM—How did it get here and where will it go. Science 361, 876–880 (2018).

2. Nogales, E. & Scheres, S. H. W. Cryo-EM: A Unique Tool for the Visualization of Macromolecular Complexity. Mol Cell 58, 677–689 (2015).

3. Lander, G. C. et al. Complete subunit architecture of the proteasome regulatory particle. Nature 482, 186–191 (2012).

4. Yan, C., Wan, R., Bai, R., Huang, G. & Shi, Y. Structure of a yeast activated spliceosome at 3.5 Å resolution. Science 353, 904–911 (2016).

5. Khoshouei, M., Radjainia, M., Baumeister, W. & Danev, R. Cryo-EM structure of haemoglobin at 3.2 Å determined with the Volta phase plate. Nat Commun 8, 16099 (2017).

6. Bartesaghi, A. et al. 2.2 Å resolution cryo-EM structure of bgalactosidase in complex with a cell-permeant inhibitor. Science 348, 1147–1151 (2015).

7. Bartesaghi, A. et al. Atomic Resolution Cryo-EM Structure of b-Galactosidase. Structure 26, 848–856.e3 (2018).

8. Tan, Y. Z. et al. Sub-2Å Ewald curvature corrected structure of an AAV2 capsid variant. Nat Commun 9, 3628 (2018).

9. Henderson, R. The potential and limitations of neutrons, electrons and X-rays for atomic resolution microscopy of unstained biological molecules. Q. Rev. Biophys. 28, 171–193 (1995).

10. Merk, A. et al. Breaking Cryo-EM Resolution Barriers to Facilitate Drug Discovery. Cell 165, 1698–1707 (2016).

11. Fan, X. et al. Single particle cryo-EM reconstruction of 52 kDa streptavidin at 3.2 Angstrom resolution. (2018). doi:10.1101/457861

12. Herzik, M. A., Wu, M. & Lander, G. C. Achieving better-than-3-Å resolution by single-particle cryo-EM at 200 keV. Nat Methods 14, 1075–1078 (2017).

13. Suloway, C. et al. Automated molecular microscopy: the new Leginon system. J Struct Biol 151, 41–60 (2005).

14. Herzik Mark A J., Wu, M. & Lander, G. C. Setting up the Talos Arctica electron microscope and Gatan K2 direct detector for high-resolution cryogenic single-particle data acquisition. (2017). doi:10.1038/protex.2017.108

15. Eklund, H. et al. The structure of horse liver alcohol dehydrogenase. FEBS letters 44, 200–204 (1974).

16. Meijers, R. et al. Structural evidence for a ligand coordination switch in liver alcohol dehydrogenase. Biochemistry 46, 5446–5454 (2007).

17. Cheng, Y., Grigorieff, N., Penczek, P. A. & Walz, T. A primer to single-particle cryo-electron microscopy. Cell 161, 438–449 (2015).

18. Scheres, S. H. W. RELION: implementation of a Bayesian approach to cryo-EM structure determination. J Struct Biol 180, 519–530 (2012).

19. Kimanius, D., Forsberg, B. O., Scheres, S. H. & Lindahl, E. Accelerated cryo-EM structure determination with parallelisation using GPUs in RELION-2. Elife 5, (2016).

20. Henderson, R. et al. Outcome of the first electron microscopy validation task force meeting. in 20, 205–214 (2012).

21. Scheres, S. H.W. & Chen, S. Prevention of overfitting in cryo-EM structure determination. Nat Methods 9, 853–854 (2012).

22. Chen, S. et al. High-resolution noise substitution to measure overfitting and validate resolution in 3D structure determination by single particle electron cryomicroscopy. Ultramicroscopy 135, 24–35 (2013).

23. Cardone, G., Heymann, J. B., Steven, A. C. Steven. One number does not fit all: Mapping local variations in resolution in cryo-EM reconstructions. J Struct Biol 184, 226–236 (2013).

24. Shibayama, N., Sugiyama, K., Tame, J. R. H. & Park, S.-Y. Capturing the hemoglobin allosteric transition in a single crystal form. J. Am. Chem. Soc. 136, 5097–5105 (2014).

25. Moore, M. J., Adams, J. A. & Taylor, S. S. Structural basis for peptide binding in protein kinase A. Role of glutamic acid 203 and tyrosine 204 in the peptide-positioning loop. Journal of Biological Chemistry 278, 10613–10618 (2003).

26. Punjani, A., Rubinstein, J. L., Fleet, D. J. & Brubaker, M. A. cryoSPARC: algorithms for rapid unsupervised cryo-EM structure determination. Nat Methods 14, 290–296 (2017).

27. Tan, Y. Z. et al. Addressing preferred specimen orientation in single-particle cryo-EM through tilting. Nat Methods 14, 793–796 (2017).

28. Zheng, J. et al. 2.2 Å refined crystal structure of the catalytic subunit of cAMP-dependent protein kinase complexed with MnATP and a peptide inhibitor. Acta Crystallogr D Biol Crystallogr 49, 362–365 (1993).

29. Dubochet, J. et al. Cryo-electron microscopy of vitrified specimens. Q. Rev. Biophys. 21, 129–228 (1988).

30. Glaeser, R. M., Typke, D., Tiemeijer, P. C., Pulokas, J. & Cheng, A. Precise beam-tilt alignment and collimation are required to minimize the phase error associated with coma in high-resolution cryo-EM. J Struct Biol 174, 1–10 (2011).

31. Lander, G. C. et al. Appion: an integrated, database-driven pipeline to facilitate EM image processing. J Struct Biol 166, 95–102 (2009).

32. Zheng, S. Q. et al. MotionCor2: anisotropic correction of beaminduced motion for improved cryo-electron microscopy. Nat Methods 14, 331–332 (2017).

33. Rohou, A. & Grigorieff, N. CTFFIND4: Fast and accurate defocus estimation from electron micrographs. J Struct Biol 192, 216–221 (2015).

34. Zhang, K. Gctf: Real-time CTF determination and correction. J Struct Biol 193, 1–12 (2016).

35. Voss, N. R., Yoshioka, C., Radermacher, M., Potter, C. S. & Carragher, B. DoG Picker and TiltPicker: Software tools to facilitate particle selection in single particle electron microscopy. J Struct Biol 166, 205–213 (2009).

36. Rosenthal, P. B. & Henderson, R. Optimal Determination of Particle Orientation, Absolute Hand, and Contrast Loss in Singleparticle Electron Cryomicroscopy. Journal of Molecular Biology 333, 721–745 (2003).

37. Herzik Mark A J., Fraser, J. S. & Lander, G. C. A Multi-model Approach to Assessing Local and Global Cryo-EM Map Quality. Structure (2018). doi:10.1016/j.str.2018.10.003

38. Wang, R. Y.-R. et al. Automated structure refinement of macromolecular assemblies from cryo-EM maps using Rosetta. Elife 5, (2016).

39. Chen, V. B. et al. MolProbity: all-atom structure validation for macromolecular crystallography. Acta Crystallogr D Biol Crystallogr 66, 12–21 (2010).

40. Adams, P. D. et al. PHENIX: a comprehensive Python-based system for macromolecular structure solution. Acta Crystallogr D Biol Crystallogr 66, 213–221 (2010).

41. Barad B.A. et al. EMRinger: side chain-directed model and map validation for 3D cryo-electron microscopy. Nat Methods 12, 943–946, doi:10.1038/nmeth.3541 (2015).

